# A Liquid Chromatography-Mass Spectrometry Method for Screening Disulfide Tethering Fragments

**DOI:** 10.1101/138941

**Authors:** Kenneth K Hallenbeck, Julia L. Davies, Connie Merron, Pierce Ogden, Eline Sijbesma, Christian Ottmann, Adam R. Renslo, Christopher Wilson, Michelle R. Arkin

## Abstract

We report the refinement of a high-throughput, liquid-chromatography/mass spectrometry (LC/MS)-based screening method for the identification of covalent small-molecule binders to proteins. Using a custom library of 1600 disulfide-capped fragments targeting surface cysteine residues, we optimize sample preparation, chromatography, and ionization conditions to maximize the reliability and flexibility of the approach. Data collection at a rate of 90 seconds per sample balances speed and reliability for sustained screening over multiple, diverse projects run over a 24-month period. The method is applicable to protein targets of various classes and a range of molecular masses. Data are processed in a custom pipeline that calculates a % bound value for each compound and detects false-positives by calculating significance of detected masses (‘signal significance’). An example pipeline has been made available through Biovia’s ScienceCloud Protocol Exchange. Data collection and analysis methods for the screening of covalent adducts of intact proteins are now fast enough to screen the largest covalent compound libraries in 1–2 days.

## Introduction

The last decade has seen an increase in the development of covalent inhibitors as potential therapeutic agents. This interest has been driven by an appreciation of the advantages of covalent mechanisms of inhibition^1–2^, including the ability to overcome resistance, such as in EGFR gatekeeping mutations^3^, the opportunity to increase affinity for otherwise ‘undruggable’ targets, and distinct pharmacokinetic properties due to very long target-residency times^4–5^. A barrier to the pursuit of such compounds has been the perception that electrophilic drugs present greater risk due to nonspecific binding to off-targets, formation of reactive metabolites, or rapid inactivation by reaction with glutathione or other endogenous nucleophiles^6–8^. However, the design and synthesis of covalent inhibitors, particularly targeting cysteine residues, has proven an effective discovery approach for select targets and therapeutic areas^9^. Furthermore, covalent inhibitors have been used as chemical probes of proteins with native or engineered cysteine residues. These success stories have utilized reversible adduct formation, such as disulfides^10^ and cyanoacrylamides^11^, or irreversible electrophiles^12–14^.

As interest in covalent drug discovery has grown, so have analytical techniques to screen for adduct formation, as well as chemical methodologies to prepare disulfide-based and electrophilic compound libraries^12, 15–16^. Despite these improvements, the largest reported screen of an electrophile library involved just 1000 compounds^13^, similar in size to our 1600-member disulfide-fragment library. Library sizes reflect several challenges inherent to the goal of discovering selective covalent inhibitors. First, adduct-forming libraries are generally custom synthesized^12, 14–16^ to normalize chemical reactivity and optimize structural diversity. Ideally, covalent ligand binding involves initial non-covalent recognition of the protein surface, followed by reaction with a proximal nucleophilic residue on the protein. If a compound is too reactive, binding is dominated by the energetics of covalent bond formation and is insensitive to molecular recognition (such chemotypes are unfortunately ubiquitous in many HTS libraries, and act as “pan assay interference compounds”, or PAINS)^17^. At the same time, small changes to compound structure can impact chemical reactivity through electronic or steric effects, obscuring underlying structure-activity relationships that derive from molecular recognition of the target. Well-designed libraries therefore seek to normalize reactivity, either by selecting electrophiles with lower functional-group sensitivity^14^ or by separating the diverse structure elements from the reactive group using linkers^10^. The design of covalent compound libraries and the development of effective covalent screening conditions must therefore control for the differing reactivity of screening compounds, and/or include counter-screens to establish selectivity^2^.

When identifying covalent ligands is the goal, it is reasonable for the primary screen to detect the formation of a covalent bond, with secondary screens for biochemical and cellular activity. Methods for measuring covalent protein modification are usually based on liquid-chromatography mass spectrometry (LC/MS), analyzing either intact protein or proteolytic peptides (LC/MS/MS). The chromatographic step in tandem MS generally takes > 10 minutes and is therefore incompatible with demands of high-throughput screening (HTS), where seconds per sample is ideal. Intact protein detection has been reported at ~3min/sample in LC formats that take advantage of Ultra-Pressure Liquid Chromatography (UPLC)^18^ and as quickly as 1.5 min/sample at high concentrations (> 10 μM) with flow-injection analysis^19^. Solid-phase extraction MS (SPE-MS) has been shown to be a viable alternative to LC/MS with reported speeds of 20s/sample^13^. While fast, SPE-MS does not allow fractionation of complex samples through chromatography. Typically, only the expected masses – rather than a full spectrum – are recorded, which can lead to false positives for noisy spectra and loss of information about multiple adduct formation.^13^ Finally, SPE-MS is a relatively insensitive MS method, using high ng/low µg amounts of protein/injection; screening is therefore done with micromolar concentrations of protein, limiting the ability to distinguish high-affinity binders and measure apparent binding affinities.

Here, we report an intact protein LC/MS method for the rapid (84 sec/sample) screening of covalent small molecules using a custom 1600 compound library of disulfide–bearing fragments. While 4-fold slower than available SPE-MS methods^13^, our approach takes advantage of efficient UPLC desalting to inject less sample. For an example 19.5 kDa protein, Campuzano and colleagues’ SPE-MS method has detection limits of 40 ng and screening injections of 400 ng (10 μL of 2 μM)^13^. Across 31 proteins of various molecular weights (MW), our method has detection limits of 0.2–20 ng, with screening injections of 12–120 ng (6μL of 100–500 nM). This enables screening of our library, including assay development, with as little as 20 µg of purified protein. We have applied the method to a range of protein classes, collecting high-quality spectra at a speed capable of sustainably screening 1000 compounds/day. Custom pipelines facilitate data processing and analysis. Since this method uses commonly available equipment, has low protein consumption, and is analyzed with publicly available computational tools, it can be readily adopted in other laboratories.

## Materials and Methods

### Protein Expression and Purification

Desired WT sequences of target proteins were cloned from their respective cDNA into a pET15b plasmid containing a 6xHis affinity tag followed by a TEV protease cleavage site at the N-terminus. Cysteine mutations were made via Megawhop PCR^20^ or QuikChange^TM^ Site-Directed Mutagenesis Kit (Agilent). All constructs were verified by DNA sequencing.

Recombinant protein expression protocols for targets in Table 1 varied to obtain optimal yield. For example, Lfa1, Mac1 and 14-3-3σ were grown in *E. coli* Rosetta 2(DE3) at 37 °C until OD600 reached 0.3. The temperature was reduced to 25 °C and at OD600 = 0.6 expression was induced with 0.25 mM IPTG followed by overnight culture. Cells were harvested by centrifugation, resuspended in 50 mM HEPES pH 7.5, 500 mM NaCl, 10 mM MgCl, 0.25 mM TCEP, 10 mM imidazole and 5% w/v glycerol, and lysed by microfluidization (Microfluidics). The soluble lysate fraction was incubated with HisPur^TM^ Cobalt resin (Thermo), washed and eluted by gravity flow in lysis buffer containing 150 mM imidazole. To remove the 6xHis affinity tag, purified protein was incubated overnight at 4 °C with 0.5 mg recombinant TEV protease with its own 6xHis affinity tag and dialyzed with an excess of 20 mM HEPES pH 7.5, 250 mM NaCl, 10 mM MgCl, 0.25 mM TCEP and 5% w/v glycerol. TEV protease and uncleaved protein were removed by re-pass over a HisPur^TM^ Cobalt resin column equilibrated in lysis buffer. Cleaved and re-passed protein was further purified by size exclusion chromatography on a Superdex 75 16/600 column (GE Healthcare) in 20 mM HEPES pH 7.5, 250 mM NaCl, 10 mM MgCl, and 5% w/v glycerol. Protein purity was confirmed via SDS-PAGE. WT protein identity and cysteine mutation presence were confirmed by intact protein LC/MS on a Xevo G2-S (Waters). Pure protein was concentrated to >5 mg/mL, flash frozen in LN2 and stored at −80 °C.

**Table 1.**
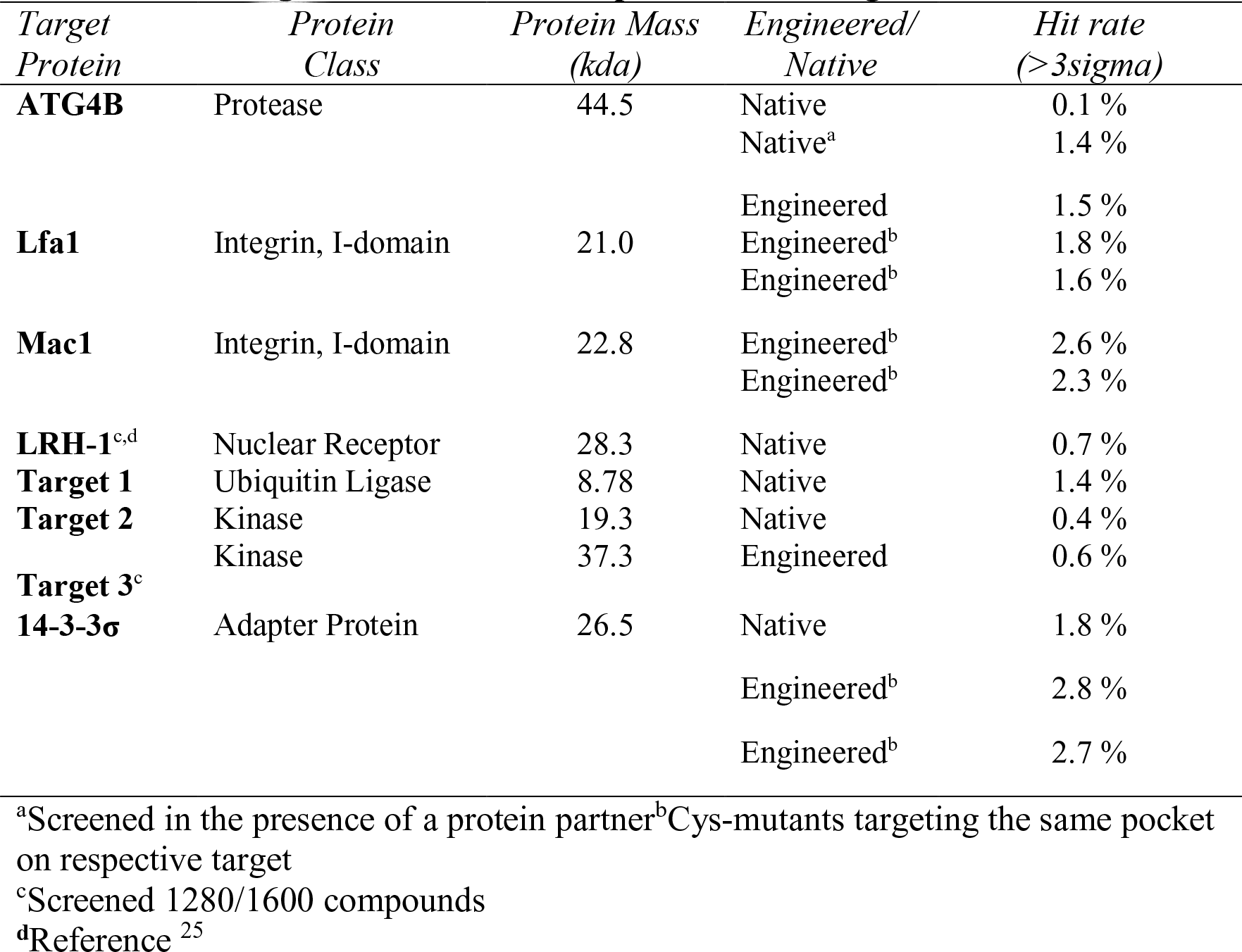
Screening Outcomes Across Representative Targets.

### Compound Library

A custom library of 1600 disulfide exchangeable compounds available at the UCSF Small Molecule Discovery Center (SMDC) was synthesized using parallel methods as previously described^15–16^. For screening, the compounds were arrayed in 384w plates as 50 mM solutions in DMSO.

### Disulfide Tethering

Protein constructs containing target cysteines were diluted to screening concentration (Table 1) in 20 mM Tris pH 8.0. 15 μL of the dilute protein was plated into columns 3–22 of a 384-well Low Volume V-Well Greiner Bio plate, with water in rows 1–2 and 23–24. 30 nL of disulfide-capped fragments were pinned into the 320 wells containing protein with a Biomek FX (Beckman), and the reaction mixture was incubated for 1–3 hours at RT (depending on experimental determination of time-to-equilibrium). Two plates of compounds were prepared simultaneously for overnight data collection.

### Liquid Chromatography

UPLC used an I-Class Acquity UPLC (Waters) using a BEH C4, 300 Å, 1.7 µm x 2.1 mm x 50 mm column. A flow rate of 0.4 mL/min was used with the gradient scheme outlined in SI Fig 1, operating at pressures 8000–10,000 psi. Mobile phase A was H_2_O + 0.5% formic acid and B was acetonitrile + 0.5% formic acid. 6 μL of sample was drawn from 384-well low-volume plates and injected, a 12 s process. Post-injection wash of 50:50 MeOH:H2O added 6 s to yield a total experiment time of 84 s. The UPLC was diverted to waste from time = 0 to 0.30 min, and again after 0.90 min; eluent from 0.30 to 0.90 min was routed to the mass spectrometer for detection. UV absorbance at 280 nM was collected for troubleshooting purposes during the experiment time of 0.30 min to 0.90 min.

**Figure 1.**
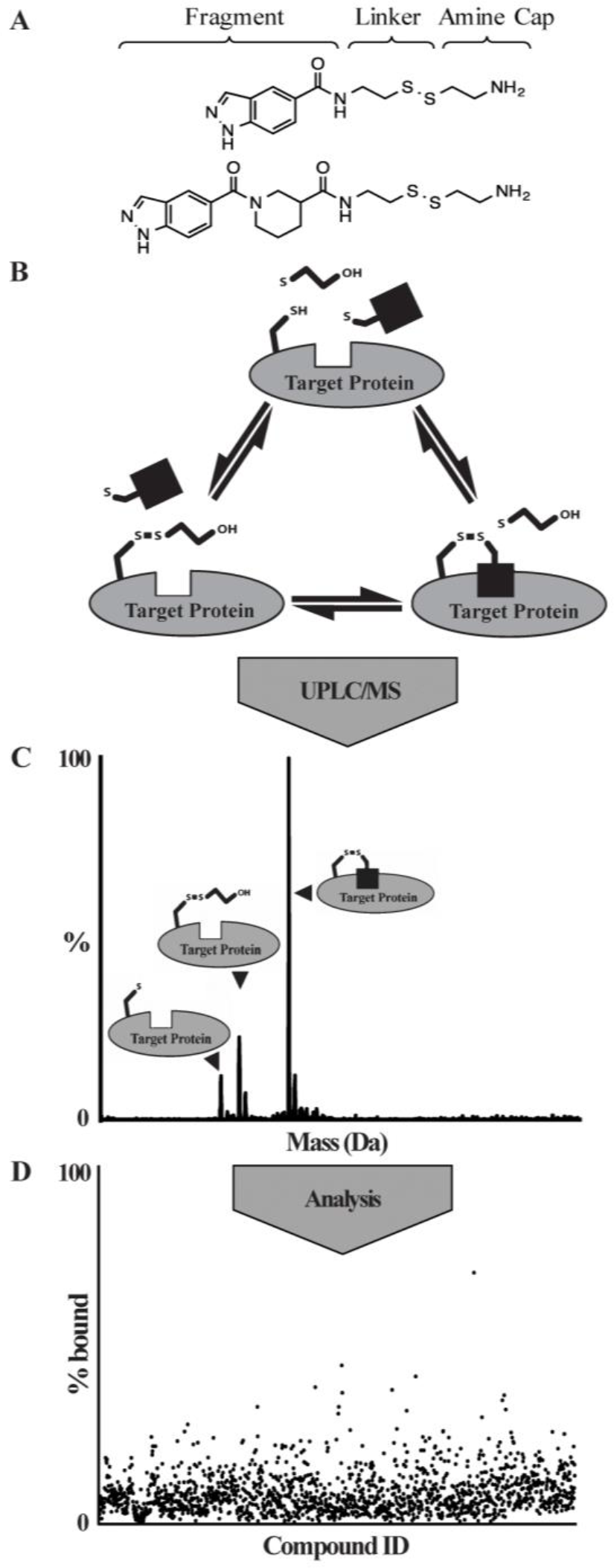
LC/MS Screening Workflow. A) Examples of structures from Tethering library^15–16^. B) Labeling reaction scheme. Target protein, βME, and various fragments (black square) are mixed in individual wells of a 384 well plate and incubated until equilibrium. C) Rapid UPLC desalting, TOF detection and m/z deconvolution identifies unlabeled, βME capped, and fragment-bound protein species. D) Detected species are checked for expected fragment adduct formation and plotted as a % of protein that is fragment-bound. Results are checked for data quality and uploaded to an internal database where selection of hits for follow-up.

### Mass Spectrometry

Mass Spectrometry data was acquired on a Xevo G2-XS Quadropole Time of Flight mass spectrometer with a ZSpray ion source (Waters). Electrospray ionization (ESI) conditions were optimized for m/z signal intensity of a Leucine Enkacephalin dimer (LeuEnk) (Waters) peak at 1111.6 Da by direct infusion of 200 pg/µL solution MeOH:H2O with 0.1% FA. The dimer peak was used because it falls in the typical m/z range of analyzed protein charge envelopes (1000–2000 Da). Two ng/µL LeuEnk was additionally used as a detector control with the ZSpray LockSpray system. Screening experiments were done at a capillary voltage 3.20 kV, cone voltage 40 V, source temperature 150 °C, desolvation temperature 650 °C, cone gas 50 L/hr, desolvation gas 1200 L/hr. Data was collected at 1 spectra/second from 50–5000 m/z.

### Limit of Detection Experiments

Limit of Detection (LOD) experiments were run using the LC/MS conditions reported above. Protein samples were 2-fold serial diluted from 500-5 nM in 10mM TRIS pH 8.0 using Optima LC/MS-grade water (Fisher). Injections of increasing concentration were monitored by manual inspection of the chromatogram until a protein peak began to appear (between 0.75–0.90 min). The LOD was defined as the first concentration at which the processing parameters below yielded the expected deconvoluted mass.

### Data Processing

Raw LC/MS data files were batch processed with Waters OpenLynx within a MassLynx v4.1 environment. A maximum entropy algorithm for mass deconvolution, MaxEnt1, was used on background subtracted m/z spectra from the portion of the LC chromatogram containing protein signal. Peak picking of the chromatogram was performed with parameters noted in SI Fig 2 and always fell between 0.75–0.90 min. As noted in previous work^13^, rare peak-picking errors in noisy data can be manually inspected and combined prior to deconvolution. The OpenLynx processing parameters subtracted m/z background between 750–2000 Da, with background defined as ≤1% maximum intensity. The 750–2000 Da range for m/z subtraction and deconvolution was chosen for general application to a range of target MW, but can be varied to match a target protein charge envelope. Deconvolution was performed with a range of +/− 6000 Da around the target’s expected mass, a target resolution of 0.5 Da, with 20 iterations of MaxEnt1 (SI Fig 3). 384-well plates were batch-processed into one large .rpt file at an analysis rate of ~30s per sample.

**Figure 2.**
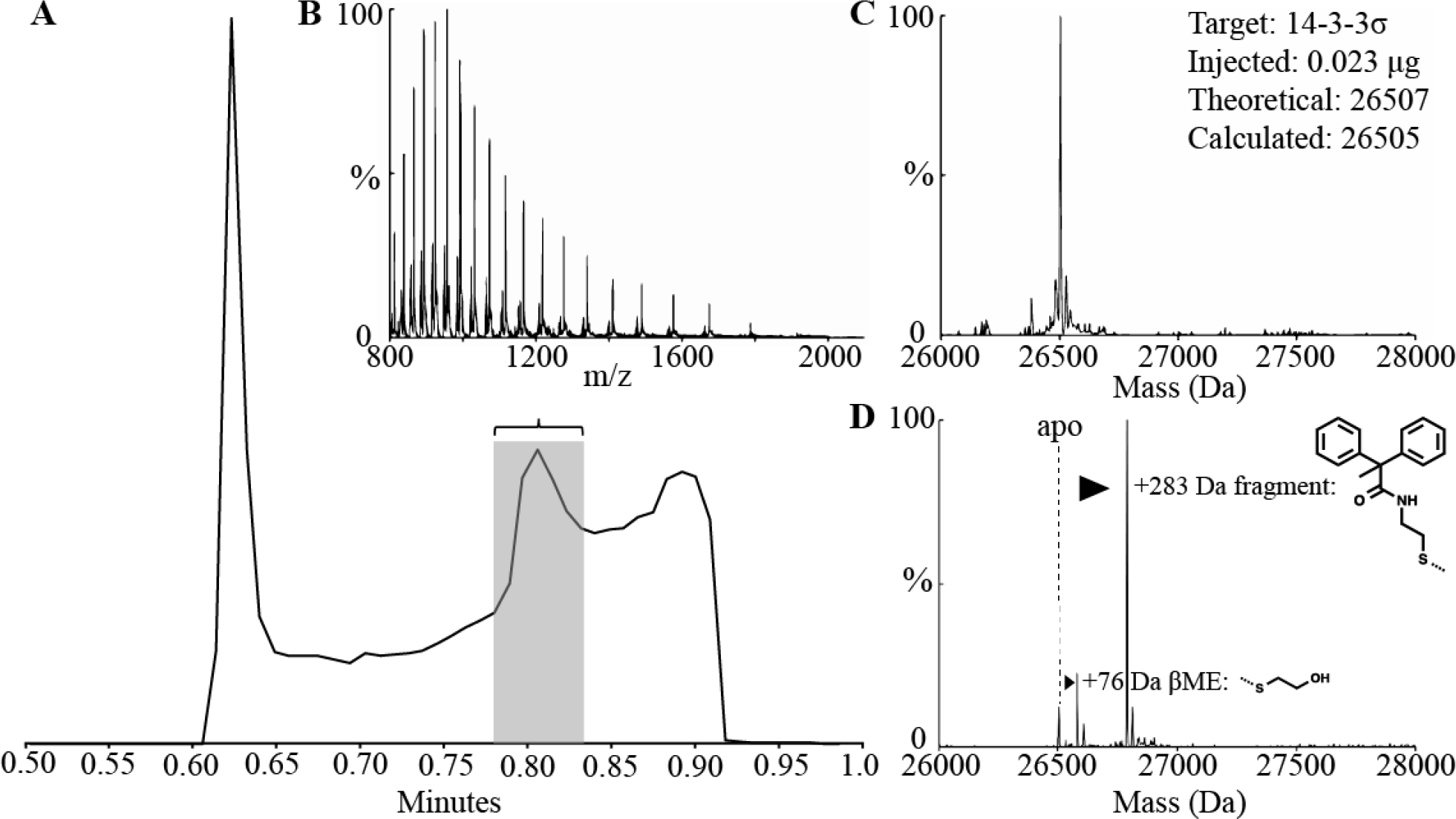
LC/MS Data and Processing. A) Total ion count trace of liquid chromatography step. Flow before 0.6 min and after 0.9 min are diverted to waste with Xevo G2S fluidics. B) The peak corresponding to protein ions (0.78–0.84 min) is combined, background subtracted, and reported as m/z. C) MaxEnt (maximum entropy) deconvolution of the m/z charge spectrum identifies the masses present in a sample containing unlabeled protein. D) MaxEnt spectrum deconvoluted from m/z shown in (B) of a reaction containing βME and screening compound, noting adduct formations.

The resulting .rpt text file was inspected for data quality within MassLynx. The expected highest abundance monoisotopic adduct masses were calculated for all compounds using Pipeline Pilot (BIOVIA) via a systematic transformation using a defined virtual reaction (SI Fig 4A). Once the expected adduct structure was verified, the highest abundance monoisotopic masses were registered using an adduct mass registration system through the Pipeline Pilot WebPort into a MySQL database. The protocol code for the adduct mass registration system has been uploaded with an example compound set to the publicly accessible ScienceCloud Protocol Exchange (Biovia) as “Adduct Highest Abundance Monoisotopic Mass Registration”. The mass of the protein-βME conjugate (cap) was calculated analogously. Protein and cap masses were registered via HiTS, a custom web application, into a MySQL database. Finally, a separate Pipeline Pilot algorithm used Eq. (1) (see Results and Discussion) to report adduct formation and Eq. (2) to provide a measure of data quality; the output was recorded in a MySQL database (SI Fig 4B).

## Results & Discussion

### Tethering screening technology

Figure 1 describes the Tethering screening methodology. Library compounds are built from structurally diverse fragment moieties (commonly < 200 Da), joined via amides, 1,2,3-triazoles, or other more extended linkers to a common aliphatic disulfide terminated with a basic amine to afford good solubility (Fig 1A). The common aliphatic disulfide moiety roughly normalizes library members’ intrinsic reactivity in disulfide exchange reactions. Fragments are mixed with proteins containing native or engineered disulfides under conditions (pH, reduction potential) that favor thiolate-disulfide exchange. Once equilibrium is reached, the reaction mixture is injected onto a UPLC/MS system; UPLC offers partial purification and ESI-TOF mass spectrometry allows determination of protein and protein+adduct masses. Sample data are provided in Figure 2.

### Method Optimization

The UPLC step was optimized for speed, signal/noise, and consistency by varying solvent flow rate (0.2–1.0 mL/min), column chemistry (C4, C8, C18), and elution strategy. A 0.4 mL/min flow over a 50 mm C4 column with a rapid (10s) gradient provided the fastest desalting which still afforded separation of proteins from post-elution noise (Fig 2A). A second ‘wash’ elution immediately followed the detected gradient to reduce carry-over of compounds and proteins on the C4 column (SI Fig 1). Flow diversion to waste before 0.3 min and after 0.9 min minimized contamination of the Xevo ion source.

We optimized the Xevo G2 LC/MS ionization conditions for detection of various proteins between 500–5000 m/z. Varying cone voltage (80–200 V), desolvation temperature (350–650 °C), the source capillary proximity to the cone, and angle toward the cone led us to the settings described in the Materials & Methods. We then performed a limit of detection (LOD) test on a series of proteins with varying molecular weight, without modifying the experimental or analysis parameters (Fig 3A). LOD was defined as the lowest concentration at which a given sample could be successfully processed in the data analysis pipeline; LOD values varied from 5–10 nM (ca. 1–5 ng per 6 μl injection; 12 proteins) to 250 nM (5 proteins). Representative chromatograms, m/z spectra and deconvoluted masses from a range of protein classes and MW are shown in Fig 3B-E.

**Figure 3.**
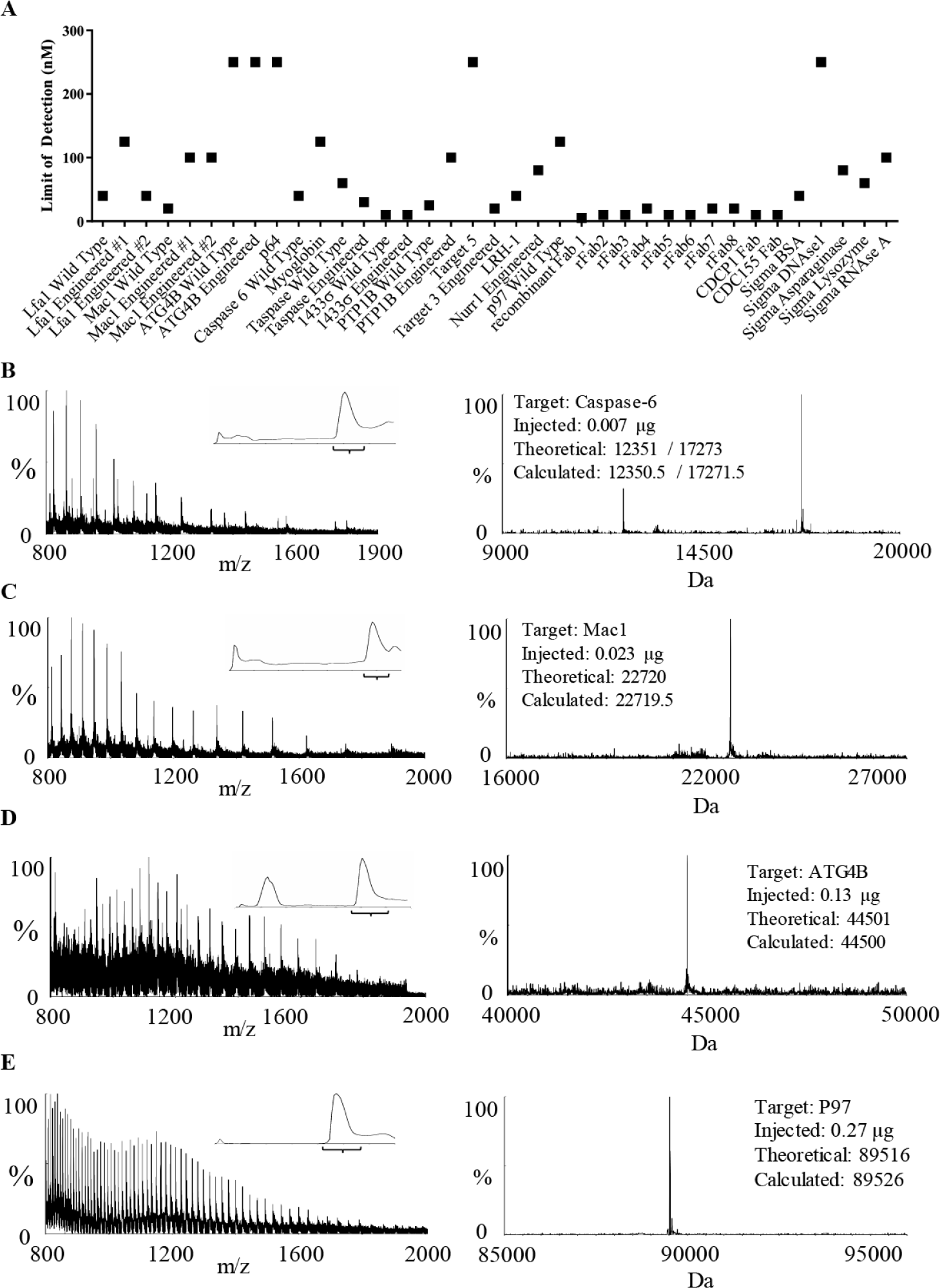
LC/MS Data Across MW and Class. A) Limit of Detection Studies. Using a 5 μL injection for a wide range of protein samples, the limit of detection ranged from 5–250 nM for our recombinant samples and a suite of controls (Sigma). B) A low MW target, caspase-6, is a tetramer containing small and large subunits, which are resolved by MaxEnt1. C) An intermediate MW target, the I-domain of Mac1 is a 22.7 KDa monomeric ligand-binding domain. D) A higher MW target, ATG4B, is a monomeric cysteine protease. E) A high MW target, P97, is a hexameric AAA+ ATPase that ionizes as a 89.5 kDa monomer. For each example protein, the left panel m/z spectrum is combined from the inset LC chromatogram and the corresponding MaxEnt1 deconvolution is shown in the right panel, along with the amount injected, theoretical and calculated mass.

### Assay Development

Assay development for screens followed a 3-step process. First, protein concentration was selected to be 2-fold LOD. For example, various cysteine mutants of adapter protein 14-3-3σ have detection limits of 10–50 nM (0.2–2.5 ng; Fig 3), and we selected a screening concentration of 100 nM. Second, tethering constructs were probed for reactivity with a titration of β-mercaptoethanol (βME), a thiol capable of forming a disulfide with an available cysteine thiolate, to confirm solvent accessibility and chemical reactivity of the target cysteine^10^. Screens were run from 100–1000 μM βME, and screening conditions were selected where a minor βME peak (ca. 20%) was present. Higher βME concentration resulted in a more stringent screen by providing competitor and increasing reduction potential of the mixture; selecting an appropriate screening concentration allowed tuning of the signal/noise and hit-rate. Notably, some cysteines showed no βME labeling during assay development but resulted in normal screening datasets.

As previously shown^10^, disulfide labeling assays are thermodynamically (vs kinetically) controlled, balancing chemical reactivity with specific small-molecule/protein interactions. One benefit of directly injecting the biochemical reaction (vs methods requiring sample pre-processing) is access to time-course and washout experiments to test labeling equilibrium and reversibility. For example, by repeated injection of 2 µL from the same well containing a 100 µL reaction of 100 µM compound, 100 nM 14-3-3σ and buffer, the reaction reached equilibrium in 5 minutes and remained stable for 45 minutes (SI Fig 5). An aliquot was then diluted 1000-fold with reaction buffer and assayed to confirm reversibility. Typically, the stability of the target at selected protein and βME concentration was tested by incubation at room temperature for 1–3 hours before analysis. A time was selected where the signal intensity was stable and no change in signal or % βME labeling was observed, indicating thermodynamic equilibrium.

### Primary Screening

The library of 1,600 disulfide fragments was stored in 384-well format in DMSO at 50 mM. 30 nL of the compound library was pinned into a reaction mixture of protein diluted into 20 mM TRIS or Ammonium Acetate pH ≥8.0, the high pH chosen to increase the concentration of thiolate and therefore facilitate thiolate/disulfide exchange. The exchange reaction was incubated until reaching equilibrium (1–3 hours) before beginning analysis.

The Acquity UPLC was equilibrated at initial conditions until ΔPSI over one minute was ≤1% of total system PSI (2–3 minutes) before beginning injections. Two plates of 320 compounds were queued simultaneously, with water in the first two and last two columns. Four dummy injections of HPLC-grade H2O were included to remove impurities in the UPLC before injections. The experiment cycle time was 84 seconds, a rate that allowed us to complete two 384-well plates overnight (15 hours) and was sustainable over long periods of use. Initial tests achieved tractable MS datasets with cycle times as fast as 50 seconds/sample but suffered from column pressure buildup and salt residue deposits on the ion source, leading us to extend cycle time to decrease maintenance requirements. The speed of the LC step relies on minimizing the amount of labeling reaction injected per sample. Injecting more than 2-fold the LOD of a target protein leads to detectable carry-over between samples. Conversely, screening too close to the LOD results in low signal/noise and increases the false positive rate from compound ion-suppression. We find that 2-fold LOD allows for rapid, sustainable LC desalting. In 24 months of operation at 84 seconds/sample, we performed 184,301 injections over 6317 hours of experimental time, consuming 134 L of mobile phase. Including idle time, regular maintenance, and intermittent instrument repair, these values translated to 8.75 hours, 251 experiments and 0.18 L of solvent per day for two years. During this time, we performed screens of several target proteins; representative screens are shown in Table 1. The method was broadly applicable and agnostic to target class or construct size. While we have not attempted to screen a protein >50 kDa, the method detected proteins ranging from 8–90 kDa (Fig 3).

### Data Processing

Raw screening data were processed with Waters OpenLynx program, software designed to apply a single Waters algorithm across large datasets. M/z data were combined across the total ion count (TIC) peak, subtracted, and analyzed with MaxEnt 1, a maximum entropy algorithm for deconvoluting intact protein mass (Fig 2, 3). These data were reported as mass vs %, in .rpt format.

Due to the volume of data and the varying quality of individual spectra, we developed a high-throughput analysis algorithm to quantify adduct formation. OpenLynx output files were read and processed using a custom Pipeline Pilot (BIOVIA) protocol to quantify binding and indicate the quality of each experiment (SI Fig 4; SI materials). Spectra were divided into small mass bins surrounding the expected masses for free protein, βME-capped protein, and protein bound to adduct, as well as one large bin for unexpected masses (SI Table 1). “Expected mass” bins included +/− 5 amu from the expected mass to accommodate resolution fluctuations due to signal/noise or drift of mass lock. The bin width could be varied from screen-to-screen to match sample quality, from +/− 2 to +/− 5 amu from target peaks. If bin overlap occurred due to a larger bin size (possible in lower quality data) or a similarity in mass between the adduct and the reductant (possible for small fragments), bins were adjusted by dividing the difference between the cap and adduct mass by 2, rounding down to the nearest integer. Within each bin, the intensities were summed and used to calculate the percent bound as in Eq. (1),

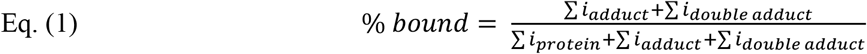

where the % of βME-protein adduct is included with ‘protein’. The protocol also checked for double-adduct formation in constructs that had alternative nucleophilic residues, e.g., two exposed cysteine residues near compound-binding sites. The algorithm additionally identified unanticipated species and adducts by reporting a maximum intensity found outside of the expected mass ranges as a secondary peak. In fact, these data were used in one study to identify and correct incorrectly drawn structures in the database.

**Figure 4.**
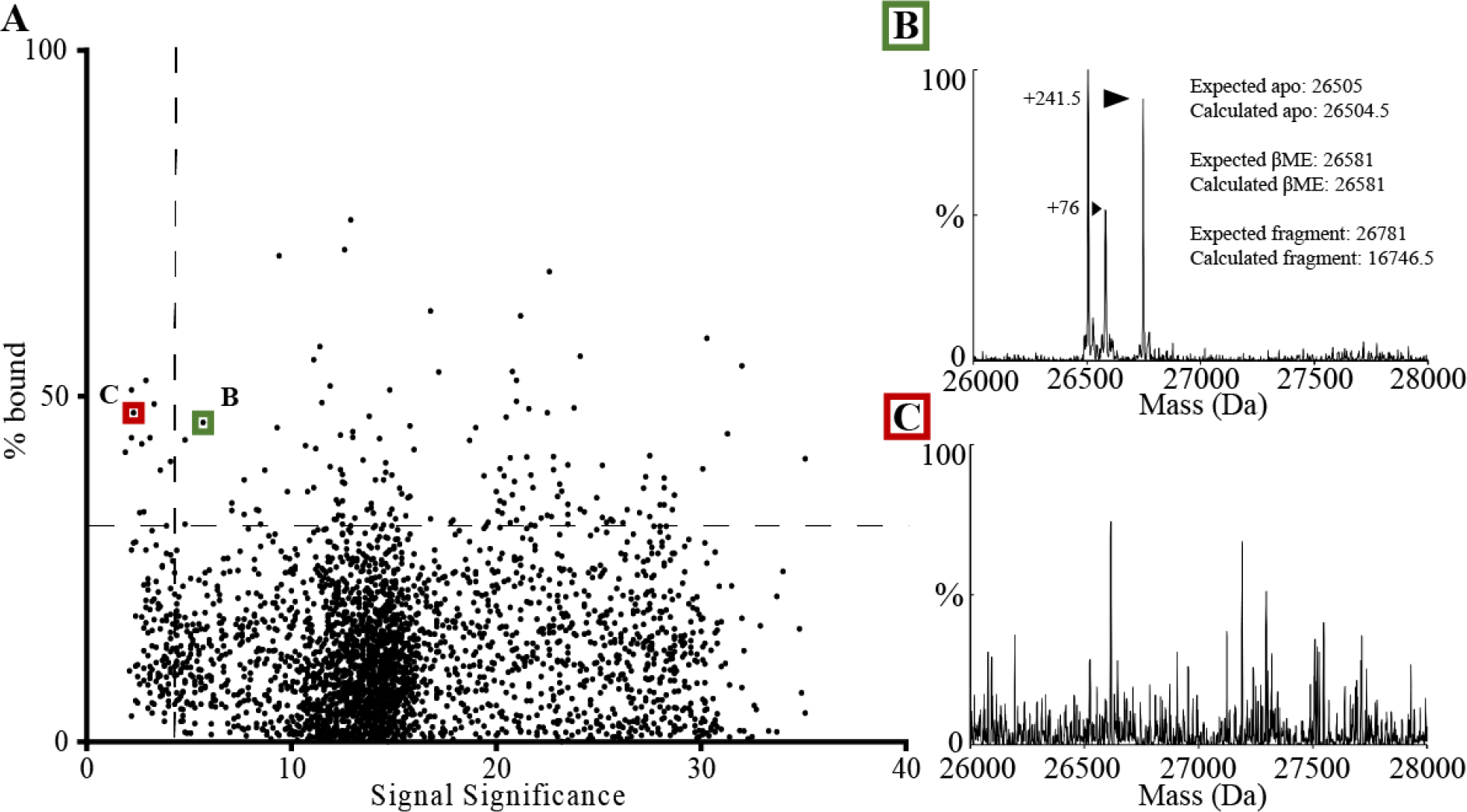
Dataset Analysis. A) A typical dataset with each of the 1600 screening compounds plotted to compare signal significance of each sample versus its calculated % bound. The horizontal dotted line is drawn at 3 standard deviations above the mean % bound. The horizontal line is drawn at an arbitrary cut-off for low quality samples determined by manual inspecting the data. B) MaxEnt spectrum for a sample (green box) with medium signal significance (<10), where adduct formation and calculated % bound are well correlated. C) MaxEnt spectrum for a sample (red box) with low signal significance (<5) where high noise has artificially inflated the % bound value.

Screening hits could be identified by plotting % bound vs compound number, e.g., as shown in Fig 1D. However, this strategy was sensitive to false positives; during the +/− 5 amu binning step, experiments with low signal/noise could report high % labeling. To provide indicators of data quality, a “signal significance number”, analogous to a signal to noise ratio, was generated by calculating the percentage of the sum of intensities in meaningful bins versus the sum of all intensities Eq. (2),

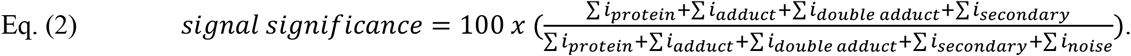

False positives with high % bound but low signal significance were readily identified by plotting the results of Eq. (1) vs. the results of Eq. (2) (Fig 4). Importantly, wells with high labeling sometimes also reported low signal significance; thus, manual inspection of hits from the lowest 5% of the signal significance range was found to be necessary (Fig 4B-C). The protocol code for the analysis step has been uploaded with an example dataset to the publicly accessible ScienceCloud Protocol Exchange (Biovia)^21^ as “Read and Analyze HTS LCMS RPT File”. Additionally, the module code is included as text in the Supplemental Information. The outputs from Eq (1) and Eq (2) were then loaded into the SMDC’s MySQL database for further analysis in a custom web application, HiTS^22^.

In conclusion, we report an optimized LC/MS method for screening intact protein for covalent adduct formation, using a library of disulfide-capped fragments. By taking advantage of advances in UPLC and ESI-TOF technology, we developed an LC method capable of more rapid (<90s) and sustainable injections than previously reported^18^. The method is capable of detecting proteins across range of molecular weights and with varying amenability to electrospray ionization (Figs 3, 4). Additionally, the labeling reaction is directly injected, facilitating kinetic studies (SI Fig 5). While our approach remains slower than extraction-based methods, it benefits from a LC desalting step to increase MS data quality, requiring less than ten ng of material per injection and 20–200 μg of protein for a full screen. The sensitivity allows the screening of low expressing and/or poorly ionizing proteins, and ability to characterize binding events over a wide affinity range. Finally, full MS spectra are collected and analyzed for unexpected adducts and for acceptable signal/noise (signal significance), allowing post-hoc inspection of data quality.

A throughput of 1000 compounds per day represents an advance in LC/MS-based screening which shifts the limiting factor in screening covalent compounds to the size of available libraries. We routinely screen and analyze our library of 1600 compounds in 3 days. Further increasing the throughput of LC/MS methods or screening compounds in mixtures will become attractive as larger libraries of electrophilic compounds become available.

Though our method is applicable to multiple target classes (Fig 3, Table 1), some targets are intractable due to protein instability at ≥10 °C or in low salt, highly reducing conditions. These limitations represent inherent facets of this approach, and targets not amenable to UPLC desalting would require a re-imagining of our screening conditions. In rare cases where the target protein is excessively hydrophobic and requires more robust chromatography, we have extended the elution gradient step from 15 seconds to 120–180 seconds, keeping all other parameters identical. While successful, the resulting screening time of 5 days could motivate the use of higher-throughput and higher-consumption methods such as SPE^13^ or Matrix-assisted laser desorption/ionization.

Applications of this and other LC/MS screening of covalent molecules extends beyond drug discovery. Adduct formation is a complex reaction, where reaction rate and equilibrium report on availability and reactivity of the nucleophile and the affinity of the probe molecule for the local environment^2^. Experiments that control for compound reactivity and affinity can probe surface ligandability^23^. Screens can be run in the presence and absence of a PPI partner or an active-site ligand to identify or confirm active-site binding or allosteric regulation^24^. Combining the control of site-directed technologies with the sampling size of high-throughput experiments generates compelling data about a target protein and the molecules that bind to it.

## Acknowledgements

The authors gratefully acknowledge Pam England, Chimno Nnadi, John Chorba, Kevan Shokat, Ambika Bhagi-Damodaran and Yinyan Tang, who provided proteins to be used in this manuscript. We also thank the National Science Foundation for a predoctoral fellowship to KKH.

